# Spatial and temporal changes in microbial communities and greenhouse gas emissions in a denitrifying woodchip bioreactor at low water temperatures

**DOI:** 10.1101/2023.04.26.538098

**Authors:** Maria Hellman, Jaanis Juhanson, Roger Herbert, Sara Hallin

## Abstract

Nitrogen (N) pollution is a major threat to ecosystems and a driver of climate change through emissions of the greenhouse gas nitrous oxide (N_2_O). Mining activities are increasingly recognized for contributing to N pollution due to undetonated, N-based explosives. A woodchip denitrifying bioreactor, installed to treat nitrate-rich leachate from waste rock dumps in northern Sweden, was monitored for two years to determine the spatial and temporal distribution of microbial communities in pore water and woodchips and their genetic potential for different N transformation processes, and how this affected the N removal capacity and possible production of undesired N species, like ammonium, nitrite and N_2_O. About 80 and 65 % of the nitrate was removed from the leachate the first and second operational year, respectively, which agreed with a decrease in dissolved organic carbon in the outlet water. There was a succession in the microbial community over time and in space along the reactor length in both pore water and woodchips, which was reflected in the genetic potential for N cycling and ultimately also reactor performance. We conclude that DNRA had minimal impact on the overall N removal efficiency due to the low relative abundance of the key gene *nrfA* involved in DNRA and the low production of ammonium. However, nitrite, ammonium, and N_2_O were formed in the bioreactor and released in the effluent water, although direct emissions of N_2_O from the surface was low. The N_2_O production in the reactor might be explained by the ratio between the genetic potential for overall denitrification and N_2_O reduction in the woodchip and pore water communities, as indicated by the low ratio between the abundance of *nir* and *nosZ* genes. Altogether, the results indicate that the denitrification pathway was temporally as well as spatially separated along the reactor length, and that unwanted reactive N species were produced at different time points and locations in the reactor. Thus, the succession of microbial communities in woodchip denitrifying bioreactors treating mining impacted water develops slowly at low temperature, which impacts the reactor performance.

## 1. Introduction

Nitrogen (N) pollution is a major threat to ecosystems and a driver of climate change through emissions of the greenhouse gas nitrous oxide (N_2_O). At the global level, there is an overall surplus of reactive N due to the wide-spread use of fertilizers and fossil-fuel combustion, resulting in increased N fluxes in the form of nitrate to aquatic ecosystems. Nitrate from mining activities caused by undetonated ammonium nitrate-based explosives is an additional source of N pollution, which is increasingly recognized as a problem. Mitigating nitrate pollution from mining activities to prevent deterioration of water quality and aquatic environments as well as climate change is a challenge due to the large volumes of water and diffuse leaching of nitrate from waste rock dumps. Fixed-bed denitrifying bioreactors, often based on woodchips, have been used for treating agricultural drainage the last decades (Schipper et al., 2010) and has recently also been employed to treat nitrate-rich water from mining activities (Nordström and Herbert, 2018) and effluents from aquaculture systems (Aalto et al., 2022; von Ahnen et al., 2019). Their low energy and maintenance requirements, together with a broad application range have promoted their use. During operation, nitrate is effectively removed through the anaerobic microbial process denitrification (Nordström and Herbert, 2018; Robertson and Cherry, 1995; Schipper and Vojvodić-Vuković, 2000) where, ideally, nitrate is converted to dinitrogen gas. However, dissimilatory reduction of nitrate to ammonium (DNRA) also removes nitrate but reduces it to ammonium, thereby retaining N in the system (Kraft et al., 2014). Further, N_2_O can be released if the denitrification process is incomplete (Philippot et al., 2011), which is the case in many denitrifying microorganisms (Graf et al., 2014). Thus, competing N-transforming processes can take place in the anoxic environment of the bioreactor, but the prevalence and importance of these processes have rarely been considered (Nordström and Herbert, 2018).

Efforts have been made to optimize process performance and bioreactor design, by considering reactor hydrology (Hoover et al., 2016; Martin et al., 2019; Nordström and Herbert, 2017; Schaefer et al., 2021), substrates for the denitrifying microorganisms (Cameron and Schipper, 2010; Hellman et al., 2021; McGuire et al., 2021; Wang and Chu, 2016) and choice of inoculum (Lefèvre et al., 2013). However, it is not until recently interest has been drawn to the microbial communities in the bioreactors. Microbial community composition of woodchip-based denitrifying systems has most often been studied during a couple of months of operation (Aalto et al., 2020; Grießmeier et al., 2017; Jéglot et al., 2021). However, few bioreactors so far have been frequently investigated for community composition over longer periods of time, despite that a denitrifying bioreactor has a life length of 10 years or more (Long et al., 2011; Robertson et al., 2008). Less is also known about the establishment of the N-transforming microbial communities in space and time in woodchip reactors, although reports on laboratory and field-scale bioreactor experiments show that the dynamics of the dominating N-transformation processes can vary over time and affect reactor performance (Hellman et al., 2021; Nordström et al., 2021).

Our aim was to determine performance of a denitrifying bioreactor in relation to development of spatial and temporal patterns in microbial communities, including genetic potential for the N-transforming processes denitrification, nitrous oxide reduction and DNRA in the pore water and woodchips during the first two years of operation. We anticipated a succession of microbial communities with a gradual development towards a complete denitrifying community (Hellman et al., 2021), both along the length of the reactor and over time. Accordingly, production of denitrification intermediates like nitrite and N_2_O were expected to decrease with distance from inlet and over time. Reactor performance included overall N removal capacity, pore water chemistry in space and time within the reactor and N_2_O and methane (CH_4_) emissions from the reactor surface. We hypothesized that overall N removal would be controlled by the degradation of woodchips, reflected by an increasing concentration of dissolved organic carbon in the reactor water DOC with time. However, depending on the ratio between nitrate and carbon substrate, competition between denitrification and DNRA can develop (Nordström et al., 2021).

## 2. Material and methods

### 2.1. Bioreactor construction and operation

The denitrifying woodchip bioreactor was built sub-surface at the Kiruna iron ore mine located in northern Sweden (67°51′ N, 20°13′ E) in 2018. The bioreactor was constructed as a 2.1 m deep excavated oblong with trapezoidal cross section, 44 m × 7 m at the ground surface and 34 m × 2 m at the bottom, lined with an impermeable 1.5 mm thick HDPE plastic geomembrane. The trench was filled with decorticated pine woodchips to a height of 1.7 m above the bottom and inoculated with activated sewage sludge (in total 2 m^3^ of a10:1 water:sludge slurry was sprinkled on the woodchips during filling). The woodchips were covered with a 0.4 m thick layer of soil (glacial till) to prevent intrusion of oxygen into the bioreactor and the whole constructions was covered with a peat layer (1 m) for insulation.

To direct the flow to the deeper regions of the bioreactor, two vertical inner walls, extending from the surface to a depth of 1.1 m, were placed at 5 m from each end of the bioreactor. At the inlet side of the first inner wall, the woodchips layer was 2.1 m thick (no glacial till) and at the outlet side of the second inner wall, the compartment was filled with crushed rock (16-32 mm) to distribute the flow over the width of the bioreactor and prevent channeling (Fig. 1). Via a pumping well, 26 m upstream the bioreactor inlet water was pumped to the bioreactor from a subsurface water reservoir (approximately 630 m^3^, filled with 100 – 200 mm crushed rock to prevent freezing) that collected leachate from a nearby waste rock pile. The water entered the reactor through a perforated pipe, 1.6 m above the bottom of the bioreactor, and flowed by gravity until it reached the outlet compartment where it discharged through a pipe leading to an outlet monitoring chamber. The outlet monitoring chamber contained a H-flume for determining the water discharge. An FDU90 ultrasound sensor (Endress+Hauser AG, Reinach, Switzerland) registered the water depth in the H-flume and flow was calculated from calibration data.

**Figure 1.**
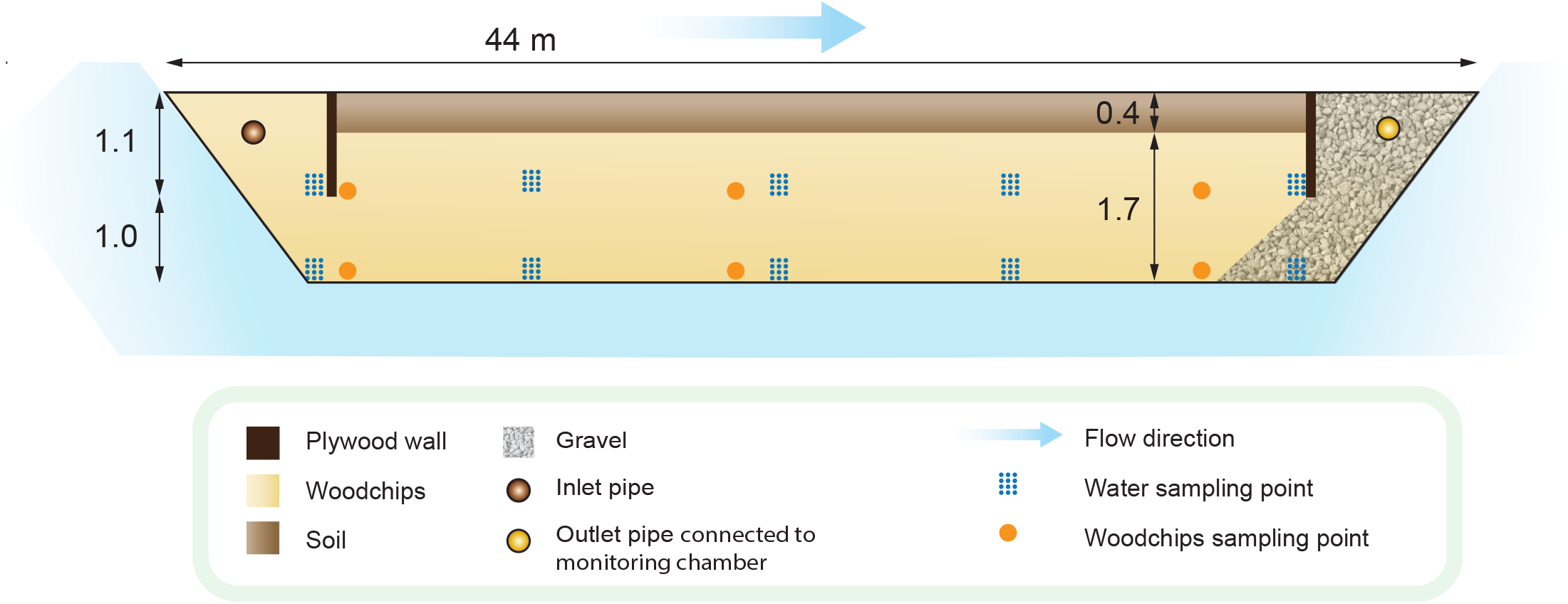
Schematic illustration of the bioreactor. The distances for the sampling points are distances from the inlet pipe. Length and depth scales are not proportional to each other. The vertical pipes and well constructions for water and woodchip sampling at the indicated depths are not shown. The insolating peat layer is not shown.

Pipes for sampling of pore water were installed at two depths, allowing for sampling at the bottom of the reactor and from 1 m above the bottom, along the length of the bioreactor, at 3.1, 11.4, 20.5, 29.2 and 37.5 m from the inlet (Fig. 1). The lower 50 cm of the pipes were screened to allow flow. Three vertical wells for sampling of woodchips (“woodchip wells”) were installed at 4.6, 19.0 and 36.3 m from the inlet in the center of the cross section of the bioreactor through the depth of the woodchip bed. The sampling wells were 30 cm in diameter and lined with a plastic net with ca 40 × 40 mm meshing for minimal disturbance of the water flow. Within the woodchip wells, nine fine mesh cylinders (2.8 × 2.8 mm meshing; 2.5 m long and 8 cm in diameter) filled with woodchips were attached to the sides. Six anchors (0.5 m diameter) for static gas chambers were installed on the reactor surface (extending ca 0.35 m below the peat surface) and were evenly distributed over the whole surface area of the bioreactor to measure fluxes of N_2_O and CH_4_. In 2019, one of the anchors was placed on the surface close to the outlet pipe in the gravel-filled outlet compartment of the bioreactor, to serve as a non-woodchip control. The anchor was moved to the bioreactor surface 2020.

Bioreactor operation started 17 September 2018, but first and second year will be used for referring to 2019 and 2020 respectively since they were full operational years. The discharge was monitored automatically via the water level in the H-flume in the outlet monitoring chamber from 28 June 2019. Tracer tests with NaBr in June 2019 and June 2020 determined the hydraulic retention time (HRT) in the reactor to ca 7.5 days at flows of 0.29 and 0.31 L s^-1^ respectively, corresponding to a bioreactor pore volume of 201 m^3^. Nitrogen in the inlet water was entirely in the form of nitrate with the average concentrations 84.1 ± 19.4 (mean ± SD; n = 17), 61.1 ± 16.6 (n = 43) and 36.9 ± 10.4 (n = 27) mg N L^-1^ during 2018, 2019 and 2020, respectively (Fig S1). The concentration of dissolved organic carbon (DOC) in the inlet water was 3.48 ±1.38 (mean ± SD, n = 16); 2.92 ±1.16 (n = 46); 5.57± 2.19 (n = 27) mg L^-1^ in 2018, 2019 and 2020, respectively (Fig S1d).

### 2.2. Sampling of water, woodchips, and gas emitted from the surface

Water for analyses of nitrate, nitrite, ammonium, and total N was collected twice a week October – December 2018, May – November 2019, and March – October 2020 from the pumping well and outlet monitoring chamber and approximately once per month in the summer periods 2019 and 2020 from the pore water sampling pipes. Water for microbial analyses was collected from the pumping well, the monitoring chamber, and the pore water pipes five times in 2019 and three times in 2020. Water from the outlet chamber was collected directly in a beaker, whereas samples from the pumping well and from the pore water pipes were collected using a peristaltic pump. Circa 2 L of water was discarded before collection of 2 L. Water for microbial analyses was filtered through a 0.22 μm pore size Sterivex ® filter until clogged, with a mean filtered volume of 1.4 L. The filters were kept at -20 °C until analysis.

For analysis of dissolved gases in the inlet (pumping well) and outlet (monitoring chamber), water was sampled by fully immersing an open 50 mL plastic syringe in the water, capping it with the piston and a stopper plug while still under water, and subsequently injecting 50 mL of water into sealed 118 mL glass bottles containing 1 mL ZnCl_2_ (50 % w/v) and 1 atm of air (2019) or N_2_ (2020). At equilibrium, 50 mL headspace gas was flushed through 22 mL vials to completely replace the air in the vials with headspace gas.

Woodchips for microbial analyses were sampled five times in 2019 and three times in 2020, within one day from the water sampling for microbial analyses, by removing one of the woodchip-filled mesh cylinders from a woodchip well and collecting six 10 cm length samples of the cylinder, representing 0-10; 10-20; 20-30; 100-110; 110-120 and 120-130 cm from the bottom of the bioreactor. The remaining woodchips were put back into the reactor bed. The samples were first kept at -20 °C and later freeze-dried.

For determining N_2_O and CH_4_ fluxes from the surface of the reactor, gas samples were collected as described in Nordström and Herbert (2018). From each of the static gas chambers nine sets of samples were collected in 2019 and three sets in 2020.

### 2.3. Chemical analyses of water and gas

Concentrations of nitrate, nitrite, and ammonium in the pore water and from the monitoring wells were photometrically determined using the Hach LKC Cuvette Test System (Hach Lange GmbH, Düsseldorf, Germany). Total N and DOC concentrations were determined in the samples from the monitoring wells using the Swedish Standard methods EN-12260:2004 and EN-1484 respectively.

Percentage of N removal (N removal efficiency) was calculated as 100 x ([total N_in_] – [total N_out_])/[total N_in_]. Nitrogen load was calculated as [total N_in_] – [total N_out_] x Q where [total N] is the concentration of total N in the water from the pumping well and outlet monitoring chamber, respectively and Q is the flow. Nitrogen removal rate was calculated as the N load divided by the bioreactor pore volume (201 m^3^).

All gas samples were analyzed for N_2_O and CH_4_ by gas chromatography (Clarus 500 GC, PerkinElmer, Waltham, MS, United States) using an electron capture- and flame ionization detector for N_2_O and CH_4_, respectively. Dissolved gas concentrations were determined via headspace equilibrium using Henry’s law and the Bunsen coefficient, without correcting for the increased pressure in the sampling bottles. The accumulated gas concentrations from the surface were recalculated to flux rates using the R package HMR (v. 1.0.1 ref).

### 2.4. DNA extraction and quantitative PCR

The Sterivex filters were detached from the filter cartridges and DNA from the water samples was extracted from the filters using the DNeasy PowerLyzer PowerSoil kit (Qiagen GmbH, Hilden, Germany). The amount of glass beads and the volumes of the reagents were modified (Supplementary methods). DNA from the woodchip samples was extracted using a combination of extraction chemistry from the DNeasy Plant Maxi kit (Qiagen GmbH, Hilden, Germany) and further purification using the Macherey-Nagel Nucleospin Soil kit (Macherey-Nagel GmbH & Co, Düren, Germany). For each sample, two separate extractions from 4 g of woodchips were combined at the end of the extraction protocol. Reagent volumes were modified (Supplementary methods).

Quantitative PCR was used to estimate the size of the total bacterial community by quantifying the 16S rRNA gene abundance (Muyzer et al., 1993). The functional genes *nirS* (Throbäck et al., 2004) and *nirK* (Henry et al., 2004), *nosZ*I (Henry et al., 2006) and *nosZII* (Jones et al., 2013), *hdh* (Schmid et al., 2008), and *nrfA* (Mohan et al., 2004; Welsh et al., 2014) were used to determine the genetic potentials for denitrification, N_2_O reduction, anammox, and DNRA. Each qPCR reaction contained 3 ng (water samples) or 1 ng (woodchip samples) template DNA, iQSYBRGreen Supermix (BioRad, Hercules, CA, United States), 15 μg Bovine Serum Albumin (BSA), and primer concentrations of 0.5–2.0 μM in a total volume of 15 μL. Two separate PCR runs were performed for each sample using the BioRad CFX or 384 Real-Time Systems. Thermal cycling conditions, primer sequences, and concentrations are available in Table S1. Standard curves were obtained using serial dilutions of linearized plasmids containing cloned fragments of the respective genes. Potential PCR inhibition was tested as described in Hellman et al. (2021) and no inhibition was detected for the DNA concentrations used.

### 2.5. Sequencing and bioinformatic analyses of 16S rRNA genes

Part of the V3-V4 region of the 16S rRNA gene was sequenced to determine the composition and diversity of the bacterial and archaeal communities in the water and the woodchips using a two-step amplification protocol. The first step was performed in duplicate 15 mL reactions containing 4 ng template DNA, 0.25 mM of primers pro515f and pro926r (Quince et al., 2011; Parada et al., 2016) with Nextera adaptor-sequences (Illumina Inc., San Diego, CA, United States), 15 μg BSA and 1 x Phusion® HighFidelity PCR Master Mix (New England Biolabs, Ipswich, MA, USA). Amplicons from the duplicate reactions were pooled and purified using Sera-Mag™ magnetic beads (GE Healthcare, Chicago, IL, USA). The second step was performed in duplicate 30 mL reactions with 10% of the final purified product from the first step as template and 0.20 mM of primers with Nextera adapter- and barcoding regions for dual labelling of the fragments. The duplicate reactions were pooled, inspected on agarose gel, purified as above, and quantified using the Qubit® fluorometer (Thermo Fisher Scientific, Waltham, MA, USA). Equal amounts of purified amplicons were pooled, the quality of the pool was checked on the Bioanalyzer and the pool of libraries was sequenced by SciLifelab in Uppsala on an Illumina MiSeq instrument using the 2 × 250 bp chemistry.

The 16S rRNA sequences were processed as described in Hellman et al. (2022), except that the non-redundant reference database SILVA version 138 was used to classify representative OTUs. SINA (Pruesse et al., 2012) was used to align nucleotide sequences of the representative OTUs to the SILVA database, and FastTree (Price et al., 2009) with the Jukes-Cantor and CAT model (Jukes and Cantor, 1969) was used to construct a phylogenetic tree from the aligned sequences. Mitochondrial and chloroplast reads were identified and removed, resulting in 10123 OTUs in the dataset. After rarefying the dataset at the depth of 10015 sequences per sample 8954 OTUs remained. For subsequent analyses, the dataset was divided into woodchips and water samples, respectively.

### 2.6. Statistical analyses

Statistical analyses were performed using R version 4.1.2 (R Development Core Team, 2021). Water chemistry, gene abundance and phylogenetic diversity data did not meet the requirements for normality, hence for comparisons between two groups Wilcoxon rank sum test was used and for comparisons between more than two groups we used Kruskal–Wallis test followed by Dunn’s test, for pairwise contrasts. Student′s t-test was used where data met the requirements for normality. Corrections for multiple comparisons were done by false discovery rate (Benjamini and Hochberg, 1995). Spearman’s rank-order correlation was used to test the relationship between DOC and N removal efficiency. Since there were no depth-related differences in nitrate concentration, functional gene abundance or phylogenetic diversity across the year per sampling location (Wilcoxon rank sum test, p > 0.05), data from the two depths were merged in the statistical analyses. For the ordinations, data from the two depths were handled separately.

Frequent OTUs were found separately for water and woodchips samples as described in Hellman et al. (2022) and Saghaï et al. (2021), resulting in 755 and 1608 core OTUs in the woodchips and water samples, respectively that retained 88.2% and 93.3% from the total OTU abundances in respective datasets. Phylogenetic diversity (Faith, 1992) of the communities was estimated using *estimate_pd* function in “btools” package. Nonmetric multidimensional scaling (NMDS) of the un-weighted Unifrac phylogenetic distances was used to visualize community patterns with “phyloseq” and “ggplot” packages, and function *envfit* in “vegan” package to correlate N cycling gene abundances with the community structure. PERMANOVA and ANOSIM (functions *adonis* and *anosim*) analyses were used to test the differences in community composition between different grouping categories. Differential abundance analyses of the core OTUs were performed using ALDEx2 (Fernandes et al., 2013) and visualized using iTOL (Letunic and Bork, 2007).

## 3. Results

### 3.1. Reactor performance and water chemistry

The total N load varied between 0.8 and 2.5 kg day^-1^. Despite a higher incoming nitrate concentration in 2019 (Fig. S1a), the load in 2020 was relatively similar due to a higher flow. In average, 79.7 ± 12.8 (mean ± SD; n = 32) and 64.7 ± 22.0 (n = 30) % of the total N was removed in 2019 and 2020, respectively and the N removal efficiency correlated with the outlet DOC concentration (Spearman′s rank correlation, p < 0.05, 2019 and 2020, both when tested separately and combined; Fig. 2). The slopes were similar despite different N removal capacities. The removal rate was significantly higher in 2019, where the average removal was 6.04 ± 2.03 compared to 4.16 ± 1.77 g N day^-1^ m^-3^ pore water in 2020 (t-test p < 0.001, n = 32 and 30, respectively). The bioreactor released similar concentrations of DOC in 2019 and 2020, but they were significantly higher in 2018 when reactor operation started than the following years (Dunn′s test; Fig. S1d).

**Figure 2.**
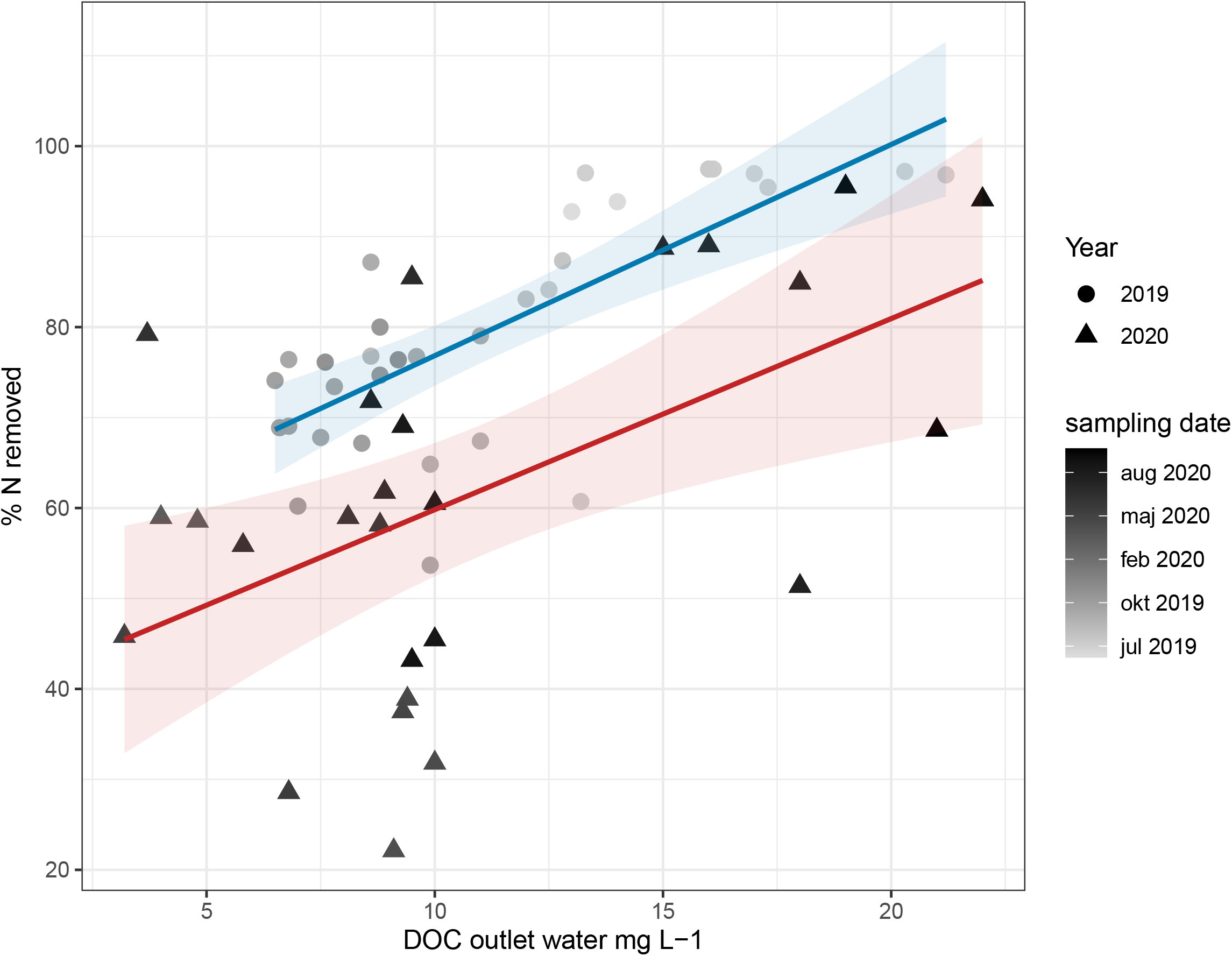
Nitrogen removal efficiency as a function of concentration of dissolved organic carbon (DOC) in the outlet water in 2019 and 2020.

In the pore water, the temporal variation of N species per sampling point was lower in 2020 compared to the first two years. Nitrate concentrations decreased with distance from the inlet, with the most rapid decrease the first year (Fig. 3a). However, nitrite was produced in the first part of the bioreactor and the change in concentration along the length of the reactor showed different patterns between the two sampling depths and years, with the highest concentrations in 2018 and 2019 (Fig. 3b and c). In contrast to the production of nitrite, ammonium production was higher towards the end of the bioreactor (Fig. 3d), with the highest concentrations in 2019 and 2020. Thus, both nitrite and ammonium were formed in the bioreactor and were also released with the outlet water (Fig. S1).

**Figure 3.**
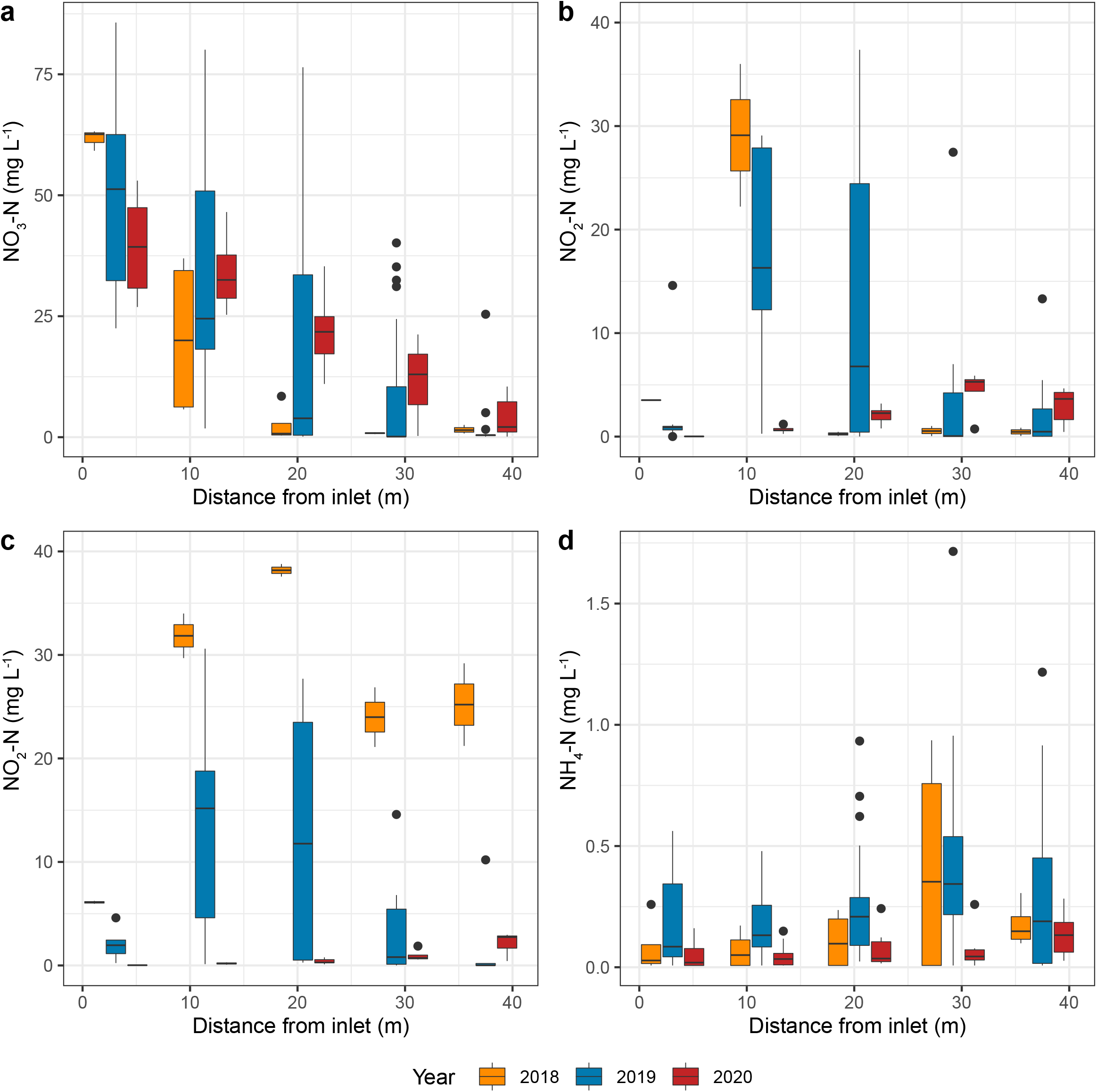
Concentration of nitrogen species in the pore water of the bioreactor 2018 - 2020. When no significant difference between layers, the concentrations shown are the average between the two layers that were sampled. a) nitrate, b) nitrite in the bottom layer, c) nitrite 1 m above the bottom, d) ammonium. Samples having concentrations below detection limit (for nitrate 0.23 mg L^-1^ and for nitrite and ammonium 0.015 mg L^-1^) are included in the plot and those samples were assigned a value of half the detection limit. Box limits represent the interquartile range with median values represented by the center line. Whiskers represent values ≤ 1.5 times the upper and lower quartiles, while points indicate values outside this range.

### 3.2. Greenhouse gases

The concentration of N_2_O in the water decreased during reactor passage in 2019, from 214 ± 61 mg m^-3^ in the inlet to 140± 123 mg m^-3^ in the outlet water (t-test p = 0.027) with the largest reductions, 97 – 98%, in June and September. By contrast, the samples from 2020, indicate net production of N_2_O in the reactor (Fig. S2a). The dissolved N_2_O-N leaving the bioreactor was in 2019 less than 0.5 % of the reactive N load and on the one occasion measured in 2020, dissolved N_2_O-N constituted 1.6 % of the N load. Emissions of N_2_O from the reactor surface were highest in the middle of the summer periods and across the whole season, there was no difference between emissions in 2019 and 2020, with mean fluxes of 9.5 and 6.6 mg N_2_O m^-2^ day^-1^ respectively, corresponding to 1.5 and 1.0 g N day^-1^ from the reactor surface area. However, when comparing between corresponding dates, the flux was significantly lower in 2020 (Fig. S2b). The N_2_O fluxes varied over the reactor surface, as did the relative contribution from each of the anchors in both years. In general, lower emissions were detected in the beginning of the reactor. The flux from the control area 2019 was low, the contribution varied between 0 and 7 % (mean 2.5 %) of the total flux.

To reveal possible short-term temporal variation in N_2_O emissions, sampling was done five consecutive days in the beginning of July 2019 (Fig. S2b). Across the week, no significant differences in flux per day over the whole surface were detected. Thus, location was more important than the time of sampling.

Methane was not detectable in the inlet water in 2019. By the small adjustment in sampling procedure, the detection limit was substantially lowered in 2020, and low concentrations of CH_4_ (0.74 ± 0.26 mg m^-3^) were found in the inlet water. After passage through the bioreactor, the levels had increased to 5 - 10 mg m^-3^, indicating production of CH_4_ in the reactor in both years. However, the fluxes of CH_4_ from the reactor surface were neglectable, and in most cases indicated a consumption rather than an emission of CH_4_.

### 3.3. Functional gene abundances and size of total bacterial communities

For the abundance of functional genes, different patterns were observed along the bioreactor between years in the water and in the woodchips (Fig. S3). In the water, with the exception for *nirK*, the abundance of genes decreased between 2019 and 2020, whereas in the woodchips an increase was observed (Fig S3a-j). The size of the total bacterial community did not change between years in the woodchips, but in the pore water there was a decrease at all sampling locations except at 42.5 m from the inlet (Wilcoxon test per sampling position; Fig S3k and l).

The ratio between clade I and clade II N_2_O reducers was below 1 in the water both years (Fig. 4a and b). In the woodchips, the ratio differed between the years, with similar ratios between woodchips and water in 2020 and overall more clade I N_2_O reducers in 2019. The ratio between genes indicating N_2_O production (sum of *nir* genes) and reduction processes (sum of *nosZ* genes) was predominantly lower than 1 in both water and woodchips, indicating a net genetic potential for N_2_O reduction (Fig 4c and d). In 2019, the ratio between the abundances of *nrfA* and sum of *nir* genes increased towards the end of the reactor in both water and woodchips, suggesting an increased importance of DNRA along the reactor, but this was not detected in 2020 (Fig 4e and f). The marker gene for anammox was not detected in the bioreactor at any location and occasion.

**Figure 4.**
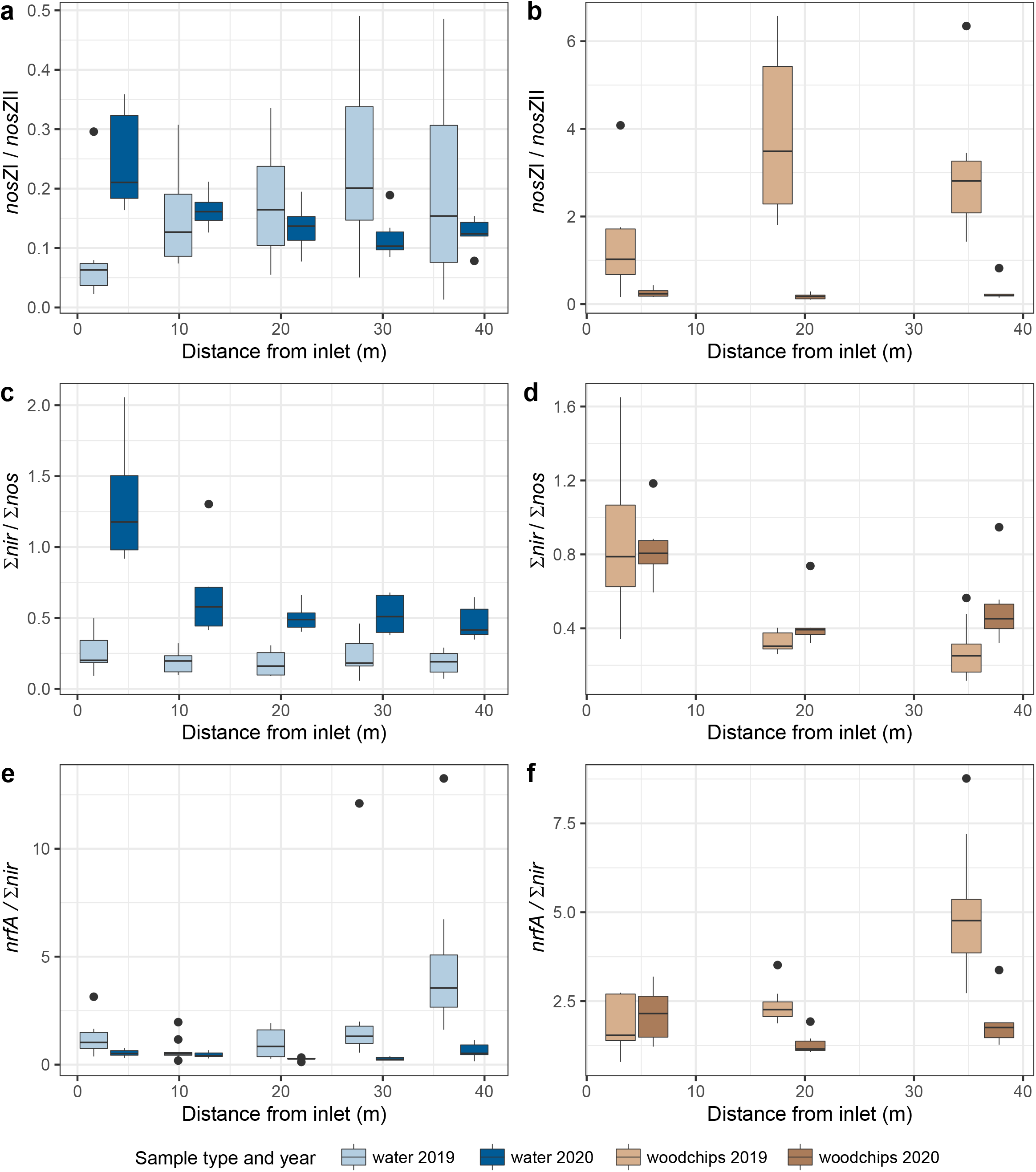
Ratios between abundance of genes along the length of the bioreactor in water (left side) and woodchip (right side). a) and b) *nosZ* clade I divided by *nosZ* clade II, c) and d) sum of *nirS* and *nirK* divided by sum of *nosZ*I and *nosZ*II e) and f) *nrfA* divided by sum of *nirS* and *nirK*. Box limits represent the inter-quartile range with median values represented by the center line. Whiskers represent values ≤ 1.5 times the upper and lower quartiles, while points indicate values outside this range.

### 3.4 Microbial community structure, diversity, and composition

The microbial community composition of frequent OTUs was different between water and woodchips samples (permanova p<0.005, anosim p<0.005). Within the water and woodchips samples, community composition was affected by year, but also by distance from the inlet, and sampling depth (permanova p<0.005, p<005 and p<0.05, respectively) (Figure 5a and b). The separation of communities between years in water samples correlated with the *nrfA/nir* gene abundance ratio, while *nir/nos* gene ratios correlated with the separation of microbial communities along the reactor (Figure 5a). In woodchips, the separation of microbial communities between years correlated with the *nosZ*I*/nosZ*II gene ratio, while *nrfA/nir* ratio correlated with the separation of communities along the reactor (Figure 5b). In addition, *nir/nos* mainly correlated with the separation of communities in the woodchips, either close to the inlet in 2019 or close to the outlet in 2020.

**Figure 5.**
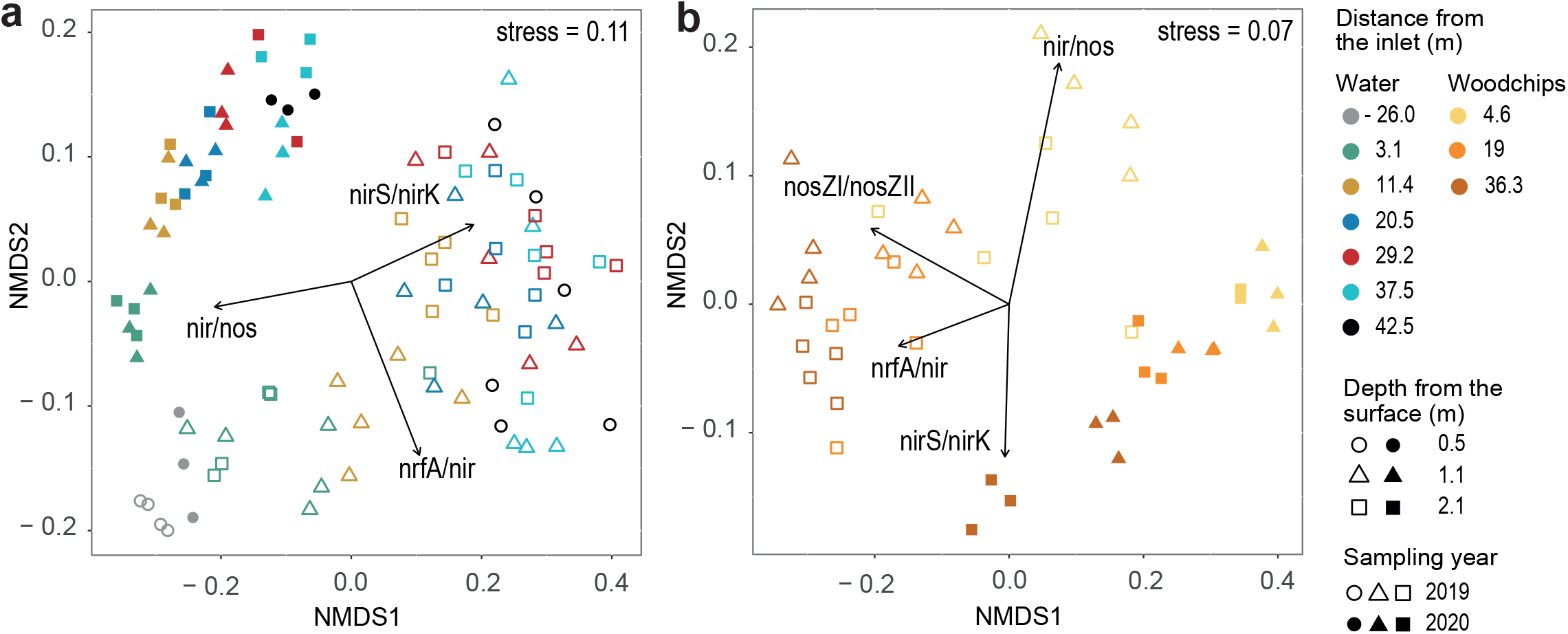
Microbial community composition in the a) water and b) woodchip samples along the bioreactor during two years of operation. For the water samples, -26.0 42.5 m from the inlet refer to pumping well water and outlet water, respectively. Ordinations are based on non-metric multidimensional scaling (NMDS) of unweighted Unifrac distances using rarefied frequent OTUs. Significant (p < 0.05) correlations between ordination axis and the abundance ratios of N cycling genes are shown as vectors which lengths are proportional to the strength of the correlations.

**Figure 6.**
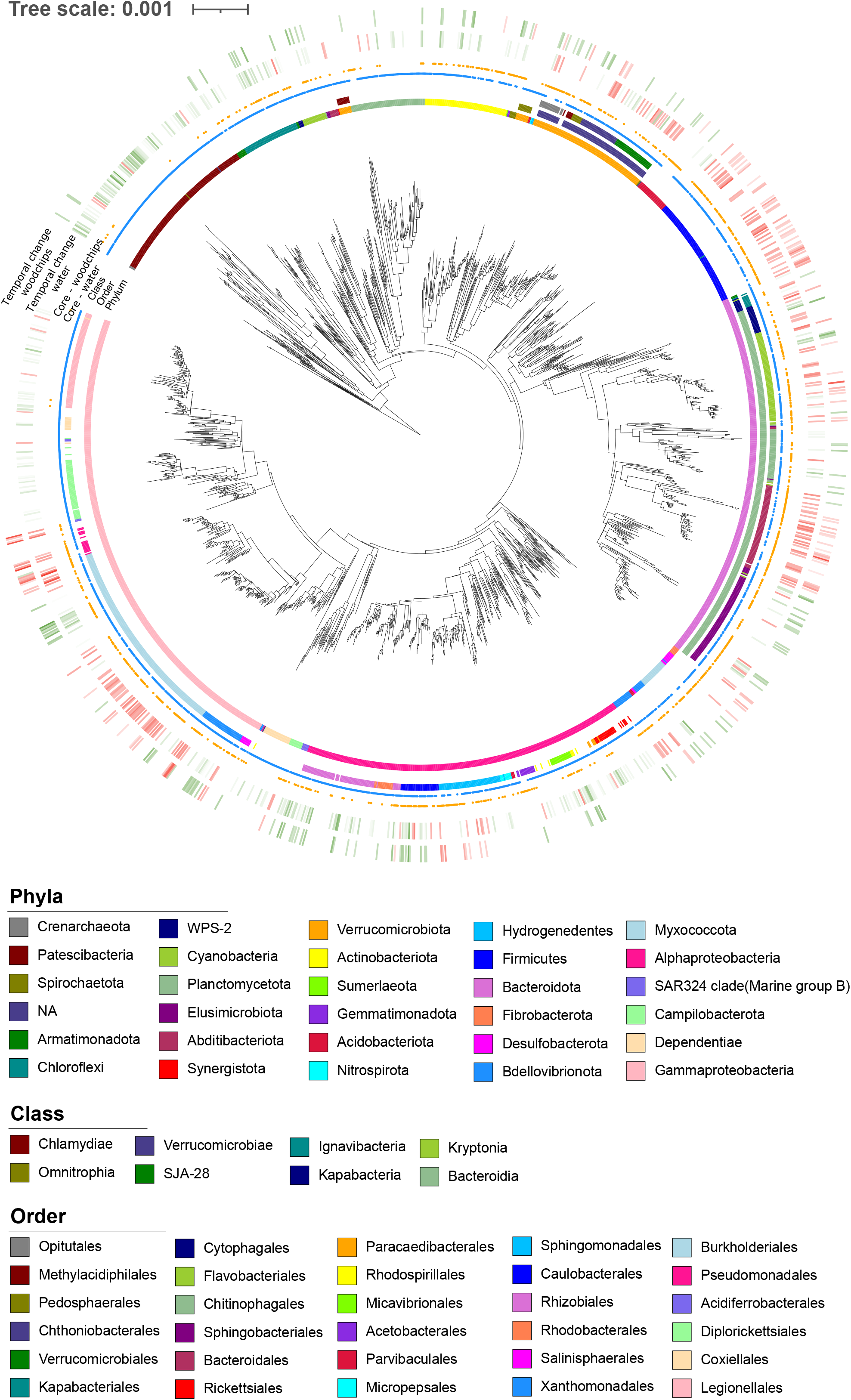
Phylogenetic tree of the core OTUs in water and woodchip samples from the denitrifying woodchip bioreactor. Different colors in the three inner rings represent taxonomy of the OTUs at the phylum, class and order levels. Blue and orange circles represent the affiliation of the OTUs in the water and in the woodchips, respectively. Red and green bars in the two outer rings represent significant temporal changes (decrease and increase, respectively) in OTU abundance in water and woodchips during the second year of operation compared to first year. Intensity of the bar color corresponds to the effect size.

The phylogenetic diversity (Faith’s PD) was in general higher in the water samples compared to the woodchips (47.2 ± 11.6, n = 95, and 33.7 ± 9.6, n = 44, respectively, Wilcoxon rank sum test p < 0.05), and higher in 2020 compared with 2019 (50.3 ± 8.4, n = 52, and 38.5 ± 12.8, n = 87, respectively, Wilcoxon rank sum test p < 0.05), except for the water from the pumping well, which demonstrated higher phylogenetic diversity than then woodchips in 2019 (66.8 ± 2.2, n = 4, and 54.4 ± 2.1, n = 3, respectively, Wilcoxon rank sum test p < 0.05). The phylogenetic diversity changed along the distance of the bioreactor (Kruskal-Wallis’ test p<0.05) and generally decreased with increasing distance in both water and woodchips samples, especially in 2019, and the decrease was fastest in the beginning of the bioreactor (Table S2). The phylogenetic diversity in the water samples from 2020 was more consistent along the distance from the inlet of the reactor, except for the samples from 3.1 m from the inlet that differed from those 29.2 and 37.5 m from the inlet pump (Dunn’s test with false discovery rate adjusted, p < 0.05). Sampling depth did not influence the phylogenetic diversity in neither the water nor the woodchips.

The microbial communities in both water and woodchips were dominated by Proteobacteria, Actinobacteriota, Bacteroidota, Firmicutes and Verrucomicrobiota, but the dynamics of their relative abundance varied along the distance from the inlet and between years (Fig. S4). In the woodchips, Actinobacteriota, Alphaproteobacteria and Verrucomicrobiota decreased along the distance from the inlet in both 2019 and 2020. However, their relative abundance were higher in 2020, while Bacteroidota and Firmicutes increased along the distance from the inlet in both years, although the relative abundance of both phyla were lower in 2020 (Figs. S4a and b, S5). The abundance of the phylum Desulfobacterota was highest in the woodchip samples most distant from the inlet (36.3 m), which was also reflected by the corresponding water sample (37.5 m). In the water samples from 2019, the dominant phyla in general followed the dynamics along the distance from the inlet in woodchips samples, but the abundances were much more consistent in the water samples 2020 (Figs. S4c and d, S5). The exception was the phylum Patescibacteria, which was in general more abundant in water samples compared to the woodchips, and also increased substantially in 2020, particularly in the middle section of the bioreactor. Unlike in woodchips, the abundance of Verrucomicrobiota was lower in the water samples in 2020 compared with 2019.

## 4. Discussion

The bioreactor removed N from the waste-rock leachate during the entire period, but with lower efficiency and removal rate the second operational year. This has been observed in other denitrifying woodchip bioreactors (Addy et al., 2016; David et al., 2016) and has been attributed to the availability of DOC (David et al., 2016; Hassanpour et al., 2017). The DOC concentrations are typically high at start-up, but decrease over time (Nordström and Herbert, 2018; Warneke et al., 2011). In this study, the decreasing N removal agrees with the observed yearly decrease in DOC in the outlet water. The start-up phase and the first year of operation was also characterized by a considerable production of nitrite in the reactor and high concentrations were also detected in the effluent, similar to what has been shown in other studies (Warneke et al., 2011; Herbert et al., 2014; Hellman et al., 2021). The nitrite production and the increase in abundance of the denitrification genes in the woodchips the second year suggest a slow development of a sufficient denitrifying community. Further, when normalizing the gene abundance to the number of 16S rRNA genes (data not shown), the percentage of denitrification genes were higher the second year, which shows that there is not just a general increase of microorganisms in the woodchips but an enrichment of denitrifiers. This pattern coincided with a higher ammonium production during the first year of operation and a higher *nrfA*/*nir* ratio, indicating that DNRA could play a role before the denitrifying community is fully developed. Nevertheless, the production and release of ammonium from the bioreactor had a negligible contribution to the total release of N. Thus, we conclude that DNRA had minimal impact on the overall N removal efficiency, which parallels the performance of a sawdust-based bioreactor treating mining impacted water but contrasts a similar reactor to the one in this study treating waste rock leachate (Nordström et al., 2021).

Nitrogen also left the bioreactor in the form of dissolved N_2_O. Both in the pore water and woodchip communities, the ratio between the genetic potential for N_2_O production and consumption (*nir/nosZ*) was well below one, which could explain the observed net N_2_O production in the reactor. The increasing *nir/nosZ* ratio in the water also offers a possible explanation for the higher concentrations of dissolved N_2_O the second year, although N_2_O was only determined once in 2020. Moreover, there was decrease in dissolved N_2_O along the length of the reactor, indicating that some of it might have been consumed by N_2_O reducing microorganisms in the reactor. The net production of N_2_O could be problematic as the dissolved N_2_O in the water could be emitted to the atmosphere downstream the bioreactor. However, direct emissions of greenhouse gases from the reactor surface were low. Less N_2_O per day left the bioreactor via emission from the surface than as dissolved in the effluent, which has also been noted from other denitrifying woodchip systems (Davis et al., 2019; Warneke et al., 2011). Since the surface of the reactor was covered with peat, the emissions of N_2_O do not necessarily reflect the production of the gas in the bioreactor compartment. Nitrous oxide reducing microorganisms in the peat layer could have used the gas produced in the bioreactor. Covering the reactor surface with soil has proven a way of mitigating N_2_O losses from the surface (Christianson et al., 2013; Manca et al., 2021). The fluxes estimated from the bioreactor are approximately 2 – 30 times higher than N_2_O emitted from fertilized agricultural soils (estimated using data from IPCC) but considering the area of a single bioreactor of these dimensions, less than 300 m^2^, the overall contribution to global N_2_O emissions would be small. Regarding CH_4_, only small amounts were produced in the reactor and the emission measurements suggests that the peat layer was a small sink of CH_4_, similar to drained peatlands (Andert et al., 2012).

The difference in the performance of the reactor between the two operational years as well as along the length of the reactor coincided with differences in the microbial community composition in both water and woodchips. The microbial community in the water samples included remarkably more OTUs compared to the woodchips, indicating that many of the microorganisms present in the water did not establish in the woodchips. The higher diversity in the water agrees with the findings by Griessmeier and colleagues (2021) in a denitrifying bed treating agricultural drainage. The higher N removal of the bioreactor in the first year may partially be explained by the higher abundance of some microbial taxa. For example, OTUs classified as orders Bacteroidales and Flavobacteriales, the phylum Firmicutes, and Gammaproteobacteria (especially OTUs related to the orders Burkholderiales and Pseudomonadales) were more abundant in 2019 compared to 2020. Most of the known microorganisms from these taxonomic groups possess genes involved in different microbial N cycling pathways (genes *nir* and *nosZ* for complete denitrification, but also *nrfA* in many members from Bacteroidota and Firmicutes). Burkholderiales are identified as key denitrifiers in woodchip bioreactors (Griessmeier et al., 2021) and also seem to play an important role in woodchip bioreactors operated under low temperatures (Jéglot et al., 2021). Burkholderiales have been found to dominate microaerobic chemostats and can play a crucial role in sustaining the dissolved oxygen concentration below the threshold of N_2_O reducers to operate (Kim et al., 2022), which offers an additional explanation to the increased N_2_O dissolved in the pore water.

The increase in the abundance of Patescibacteria in water samples the second year is interesting, since not so much is known about this phylum, although they have been found to be prevalent in aquatic environments, including oligotrophic groundwater sediment (Herrmann et al., 2019). Patescibacteria have small cells and genomes, suggesting that they may rely on host bacteria or protists. Metagenomic studies have not found pathways for chemoautotrophic metabolism, like sulfur, ammonia and nitrite oxidation (Tian et al., 2020). Instead, potential associations with abundant autotrophic taxa involved in N, sulfur and iron cycling in groundwater were identified (Herrmann et al., 2019). In addition, fermentative pathways for lactate and formate were found in Patescibacteria in a freshwater anammox column reactor and the authors suggested that Patescibacteria may provide substrate supporting growth of other bacteria in the annamox reactor (Hosokawa et al., 2021). Fermentation process have been proposed to be important to provide easily available C substrates for denitrifiers in woodchip bioreactors for efficient N removal (Nordström and Herbert, 2018), but more work is needed to confirm this, and determine the underlying mechanisms to help developing an optimal design and operation of denitrifying woodchip reactors treating N polluted water from mining activity.

There was not only a succession in the community over time, but also in space along the reactor, which was reflected in the genetic potential for N cycling, water chemistry, and ultimately also reactor performance. We further observed trends in increased denitrification gene abundance as a function of distance from the bioreactor inlet, and the gene abundance patterns largely reflected the increased nitrate reduction along the reactor, as observed previously (Herbert et al., 2014), and the decreased nitrite production as well as increased ammonium production. Depth-related differences in N transformation processes have previously been highlighted (Herbert et al., 2014), but in the present study there was only a minor effect of reactor depth on water chemistry and abundance of N-transformation guilds Altogether, this indicate that the denitrification pathway was spatially separated along the reactor and that unwanted reactive N species were produced at different locations in the reactor, especially during the first year of operation.

## Supporting information

Supplemental information Hellman et al

## Acknowledgments

This work was supported by the EIT Raw Materials (grant 17013), LKAB and the Water Joint Programming Initiative and financed via the Swedish Research Council Formas (grant 2018-02788). Sequencing was performed by the SNP&SEQ Technology Platform in Uppsala. The facility is part of the National Genomics Infrastructure (NGI), Sweden, and the Science for Life Laboratory. TheSNP&SEQ Platform is also supported by the Swedish Research Council and the Knut and Alice Wallenberg Foundation.

