## Supplemental information Hellman et al for "Spatial and temporal changes in microbial communities and greenhouse gas emissions in a denitrifying woodchip bioreactor at low water temperatures"

### Supplementary Information

#### Supplementary methods

*DNA extraction water samples*
DNA was extracted using the DNeasy PowerLyzer PowerSoil kit (Qiagen GmbH, Hilden, Germany). The Sterivex filter cartridges were opened, and the filters were detached and transferred to 7 mL screw cap polypropylene tubes. Glass beads from two bead beating tubes, 1.5 mL Power Bead Solution, and 120 µL of solution C1 was added. The tubes were run in a Precellys 24 homogenizer (Bertin Instruments, France) at 5500 rpm for 2 x 40 s with a 15 s rest between the rounds. The tubes were centrifuged for 2 min at 5500 x g. For practical reasons, each sample was divided: From each of the 7 mL tubes, 2 x 550 µL were transferred into separate 2 mL centrifuge tubes, 280 µL of solution C2 was added and the tubes were centrifuged for 1 min at 11 000 x g. The supernatant was transferred to a new tube, 260 µL of solution C3 was added and the tubes were centrifuged for 1 min at 11 000 x g. After this, the samples were combined again: The supernatants from each of the two separate tubes representing the same sample were transferred to one 15 mL tube, and 3 mL of solution C4 was added. To recover the DNA, the liquid was transferred to the kit Spin Column, arranged on a vacuum manifold. After passing the liquid through, the filter was washed with 600 µL of solution C5 and spun dry for 1 min at 11 000 x g. DNA was eluted 80 µL of solution C6. DNA concentrations in the extracts were determined using the Qubit Fluorometer (Thermo Fisher Scientific, Waltham, MA, United States).

*DNA extraction woodchip samples*
DNA was extracted using a combination of the DNeasy Plant Maxi kit (Qiagen) and the Macherey-Nagel Nucleospin Soil kit (Macherey-Nagel GmbH & Co, Hilden, Germany). 4 g of freeze-dried woodchips was re-wetted in 6 mL of sterile deionized water in a 50 mL centrifuge tube. Nine glass beads (3 mm), 0.75 mL 1.4 mm zirconium oxide beads (ca 3 g) and 2 mL 0.1 mm glass beads (ca 3 g) were added. Two replicates were made for each sample. The Plant Maxi kit was used for DNA extraction: 5 mL Buffer AP1 (preheated to 65 °C) and 10 µL RNAse was added, and the tubes were shaken in a horizontal position on a Vortex () equipped with an adaptor plate for 50 mL tubes (10 min at maximum speed and 65 °C). 1.8 mL Buffer P3 was added, and the tubes were incubated on ice for 10 min and thereafter centrifuged for 5 min at 4500 x g. The supernatant was filtered through a QIAshredder column by centrifuging for 5 min at 4500 x g and the flow-through was transferred to a new 50 mL tube. The volume was recorded and 1.5 volumes of Buffer AW1 was added and well mixed with the flow-through liquid before poured into a DNeasy Maxi spin column arranged in a vacuum manifold. The two replicates from each sample were put on the same DNeasy Maxi spin column. After passing the liquid through, the membrane was washed with 12 mL of Buffer AW2 and dried by centrifuging for 5 min at 4500 x g. DNA was eluted from the membrane in 2 x 0.75 mL Buffer AE and combined in the same tube.

The Macherey-Nagel NucleoSpin Soil kit was used to further purify the DNA extracts: Each extract was divided in two, and to 0.75 mL extract, 150 µL SL3 buffer was added. The samples were incubated on ice for 5 min and centrifuged at 11 000 x g for 1 min. The samples were filtered in a NucleoSpin Inhibitor Removal column by centrifugation. The two extracts from the same sample were loaded on the same column but the eluents were collected separately. Binding conditions were adjusted by adding 250 µL of SB buffer and the purified DNA was bound to the membrane in a NucleoSpin Soil column using a vacuum manifold. The two extracts from the same sample were combined in the same column. The membrane was washed with 500 µL SB buffer , 550 µL SW1 buffer, and 2 x 700 µL SW2 buffer. The membrane was spun dry for 2 min at 11 000 x g. DNA was eluted in 2 x 30 µL SE buffer. DNA concentrations in the extracts were determined using the Qubit Fluorometer.

**Supplementary figures and tables**

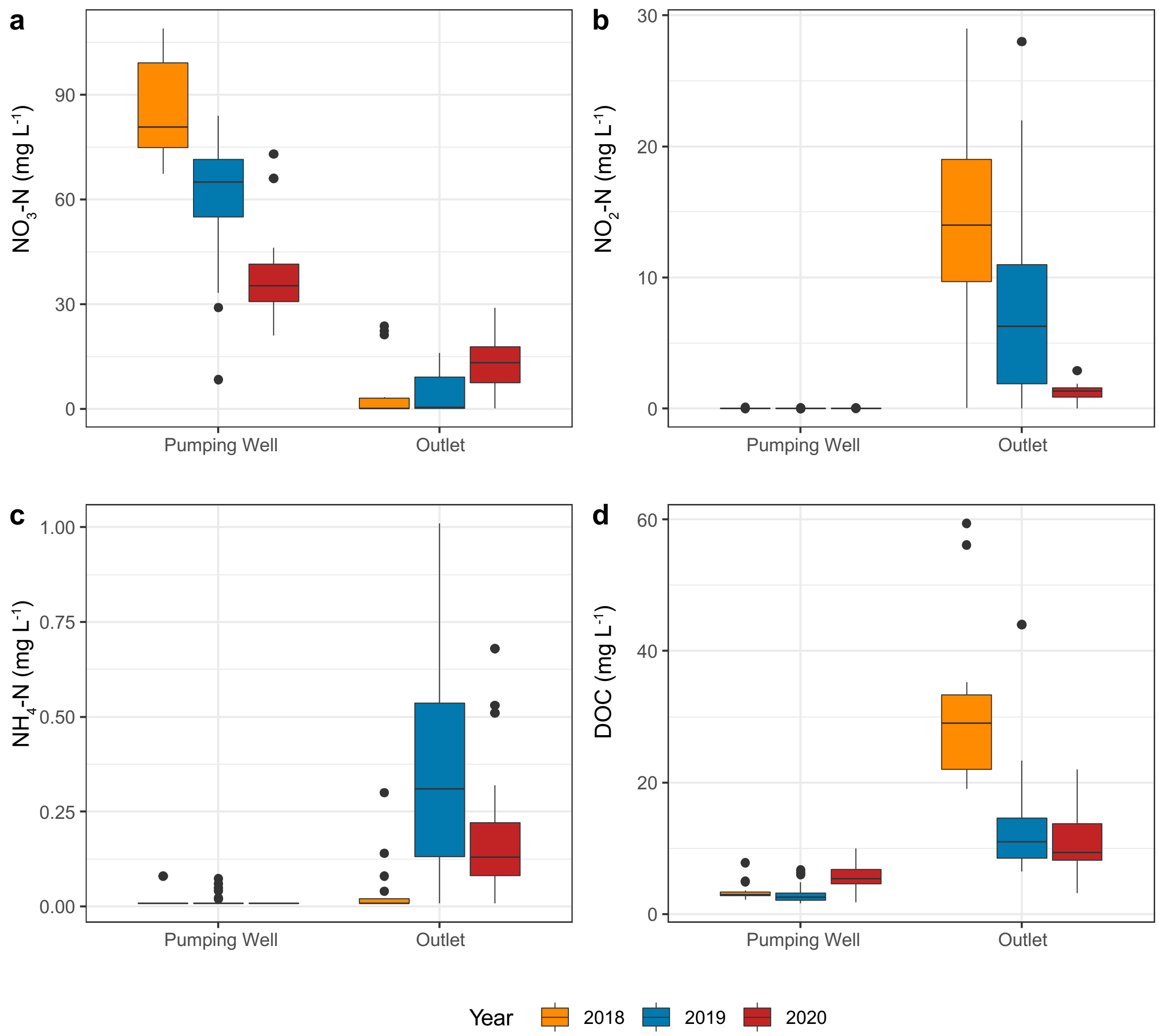


**Figure S1.** Concentration of nitrogen species and dissolved organic carbon in the pumping well with inlet water and the outlet monitoring chamber of the bioreactor 2018 - 2020. a) nitrate, b) nitrite, c) ammonium, and d) dissolved organic carbon. Samples having concentrations below detection limit (for nitrate 0.23 mg N $L^{-1}$ and for nitrite and ammonium 0.015 N mg $L^{-1}$) were assigned a value of half the detection limit. Box limits represent the inter-quartile range with median values represented by the center line. Whiskers represent values ≤ 1.5 times the upper and lower quartiles, while points indicate values outside this range.

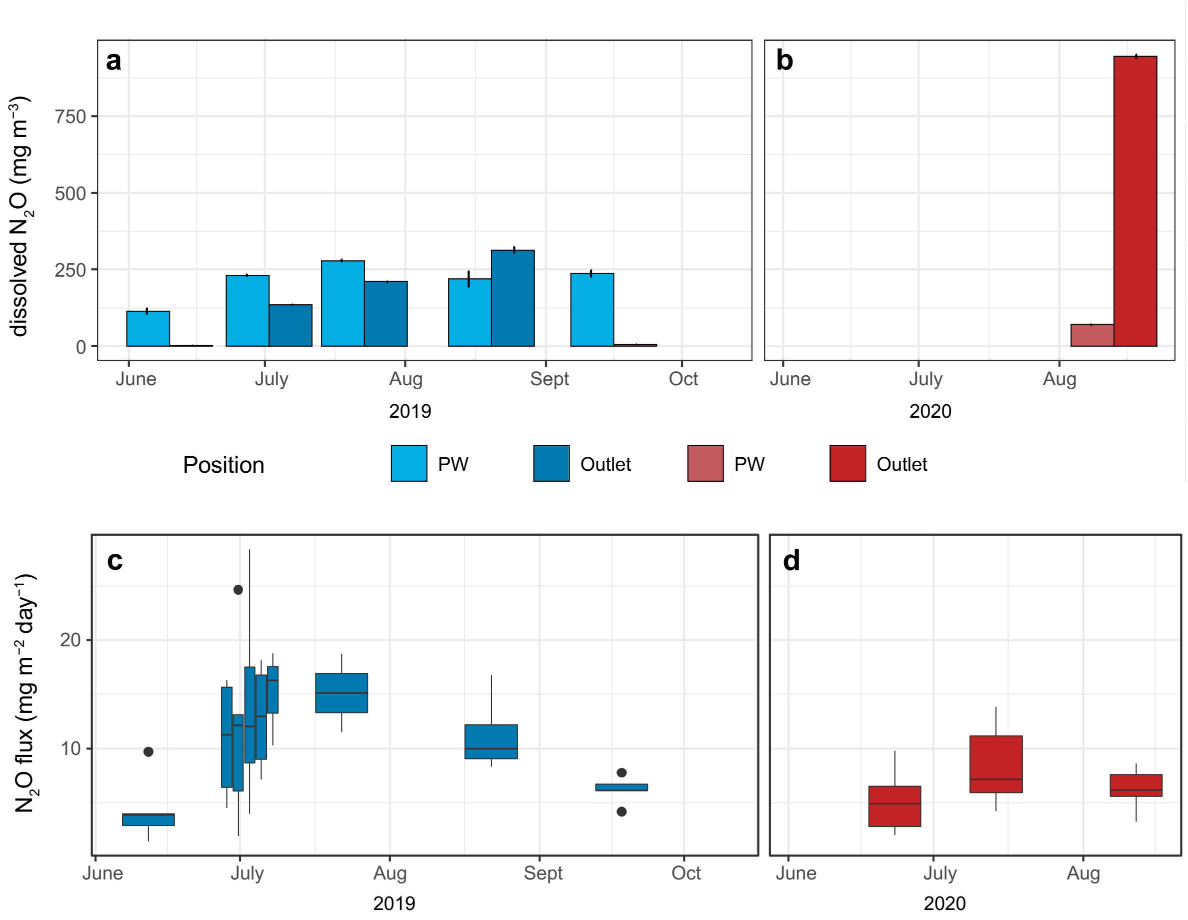


**Figure S2.** Nitrous oxide released from the reactor 2019 and 2020. a) and b) Concentrations of dissolved $N_2O$ in the pumping well (PW) with inlet water and the outlet monitoring chamber of the bioreactor. Error bars show ± 1 standard deviation, n = 3 – 4. c) and d) Fluxes of $N_2O$ from the surface of the bioreactor. Box limits represent the inter-quartile range with median values represented by the center line. Whiskers represent values ≤ 1.5 times the upper and lower quartiles, while points indicate values outside this range, n = 2 – 5, data from the control measuring point not included.

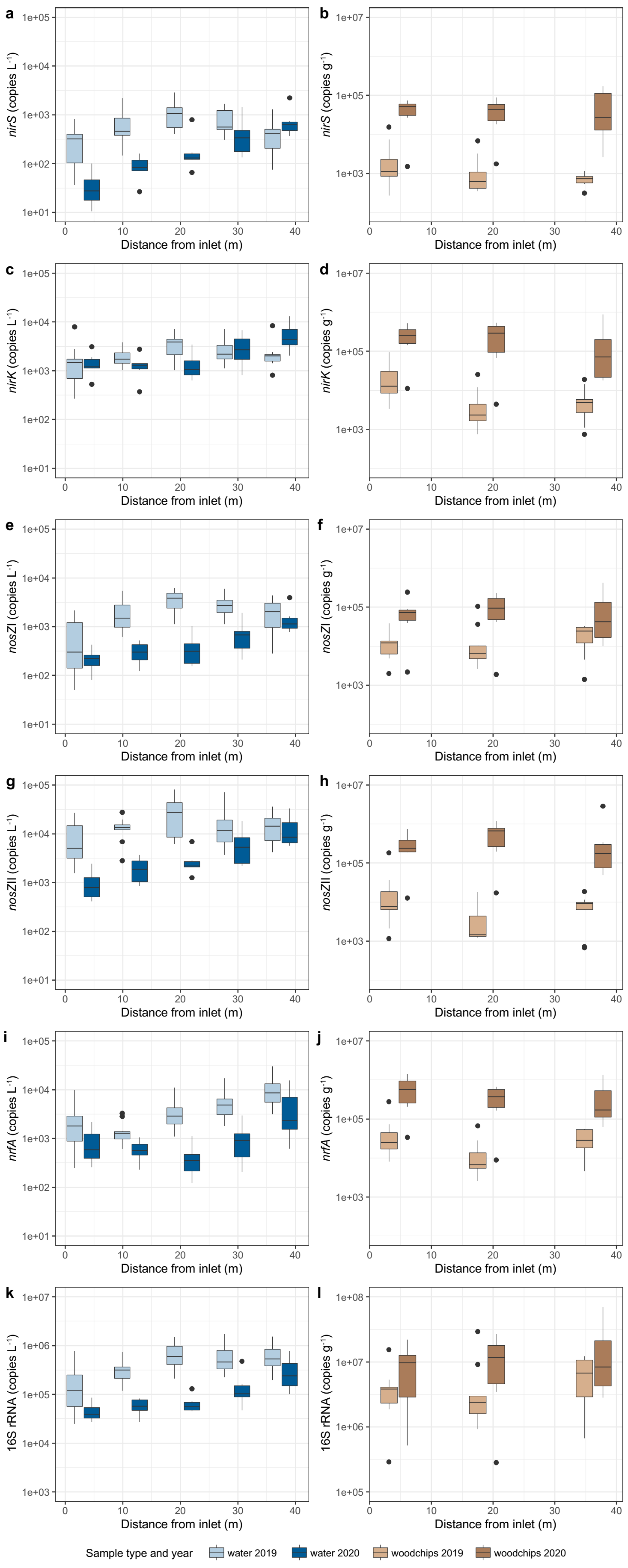


**Figure S3.** Gene abundances along the length of the bioreactor in the pore water (left side) and woodchips (right side). a) and b) *nirS*, c) and d) *nirK*, e) and f) *nosZ* clade I, g) and h) *nosZ* clade II, i and j) *nrfA*, and k and l) 16S rRNA. Box limits represent the inter-quartile range with median values represented by the center line. Whiskers represent values ≤ 1.5 times the upper and lower quartiles, while points indicate values outside this range.

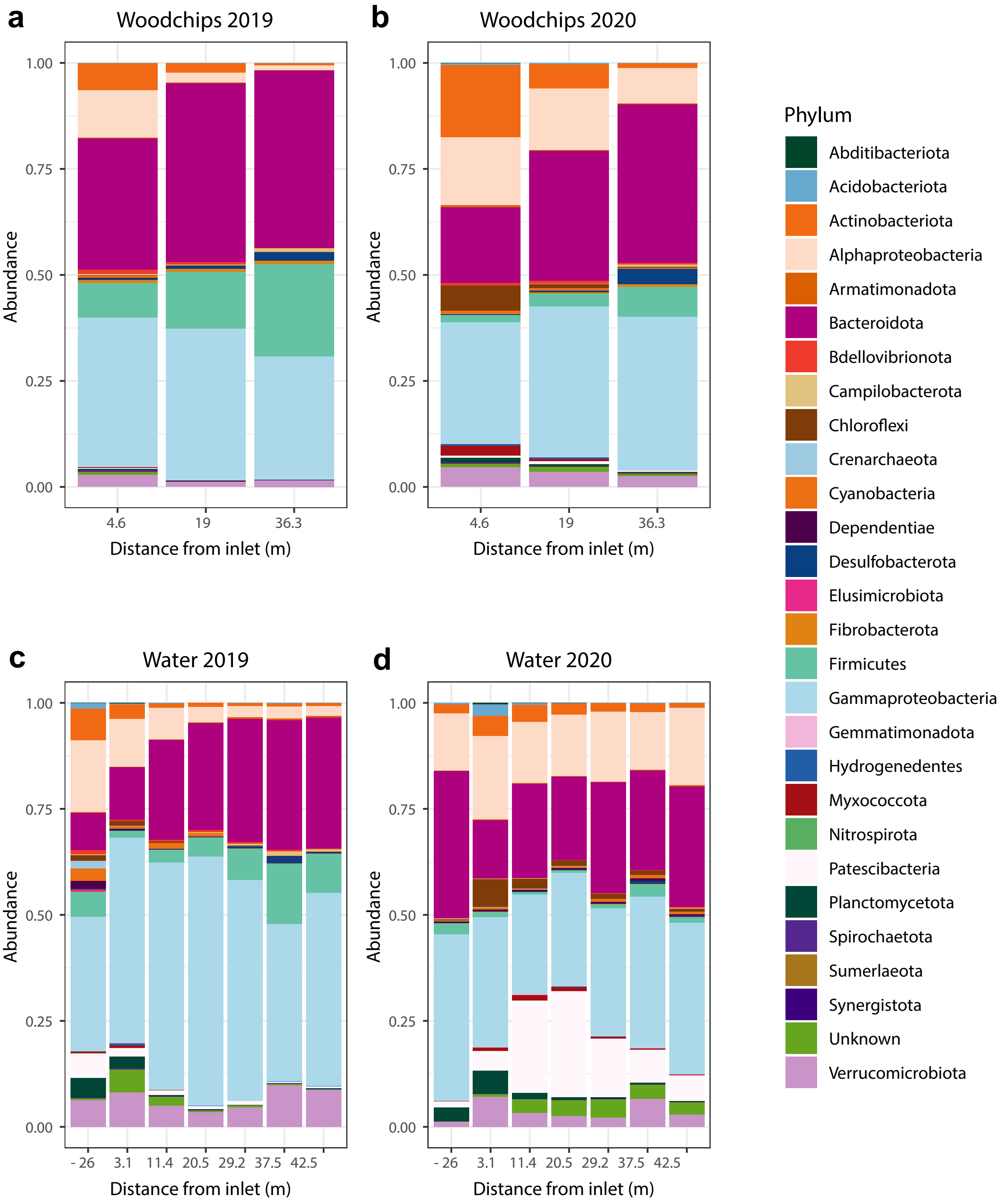


**Figure S4**. Taxonomic composition of the microbial communities in the woodchips (a and b) and water (c and d) samples along the length of the bioreactor during two years of operation. The mean relative abundances of core OTUs grouped together at the phylum level in different sample groups are shown. The phylum Proteobacteria is separated into classes Alpha- and Gammaproteobacteria.

**Table S1.** Primers and reaction conditions for qPCR.

| Gene | Primer pair | Primer sequence | Final conc | Melting | | Annealing | | Elongation | | Data acquisition | | Cycles[a] |
|---|---|---|---|---|---|---|---|---|---|---|---|---|
| (Primer ref) | | 5′- 3′ | (µM) | T (°C) | t (s) | T (°C) | t (s) | T (°C) | t (s) | T (°C) | t (s) | |
| **16S rRNA** | 341 F | CCTACGGGAGGCAGCAG | 0.50 | 95 | 15 | 60 | 30 | 72 | 30 | 78 | 5 | 35 |
| (Muyzer et al., 1993) | 534 R | ATTACCGCGGCTGCTGG**CA** | 0.50 | | | | | | | | | |
| ***nirS*** | Cd3a Fm | AACGYSAAGGARACSGG | 0.50 | 95 | 15 | 65-60[b] | 30 | 72 | 35 | 80 | 5 | 36 |
| (Throbäck et al., 2004) | R3cdm | GASTTCGGRTGSGTCTTSAYGAA | 0.50 | | | | | | | | | |
| ***nirK*** | 876F | ATCATGGTSCTGCCGCG | 0.50 | 95 | 15 | 63-58[b] | 30 | 72 | 30 | 80 | 5 | 41 |
| (Henry et al., 2004) | 1040 R | GCCTCGATCAGRTTRTGGTT | 0.50 | | | | | | | | | |
| ***nosZ*I** | 1840 F | CGCRACGGCAASAAGGTSMSSGT | 0.50 | 95 | 15 | 65-60[b] | 30 | 72 | 35 | 80 | 5 | 36 |
| (Henry et al., 2006) | 2090 R | CAKRTGCAKSGCRTGGCAGAA | 0.50 | | | | | | | | | |
| ***nosZ*II** | nosZII F | CTIGGICCIYTKCAYAC | 2.00 | 95 | 15 | 54 | 30 | 72 | 45 | 77 | 5 | 40 |
| (Jones et al., 2013) | nosZII R | GCIGARCARAAITCBGTRC | 2.00 | | | | | | | | | |
| ***nfrA*** | nrfAF2aw | CARTGYCAYGTBGARTA | 0.50 | 95 | 15 | 57-52[b] | 30 | 72 | 30 | 80 | 5 | 40 |
| (Welch et al., 2014/Mohan et al., 2004) | nrfAR1 | TWNGGCATRTGRCARTC | 0.50 | | | | | | | | | |
| ***hdh*** | Hcocl1F1 | TGYAAGACYTGYCAYTGG | 0.80 | 95 | 15 | 52.5 | 30 | 72 | 30 | 77 | 10 | 35 |
| (Schmid et al., 2008) | Hcocl1R2 | ACTCCAGATRTGCTGACC | 0.80 | | | | | | | | | |

[a]Protocols start with an activation step, 95 °C for 5 minutes and are finished with a melt curve, 15 s at 95 °C followed by 65-95 °C in 0.5 °C increments for 5 s.

[b]1 °C decrease per cycle the first six cycles

**Table S2.** Phylogenetic diversity (Faith's PD) in the water and woodchips along the denitrifying woodchip bioreactor during two years of operation.

| Sample type | Year | Distance from inlet (m) | Faith´s PD (mean ± SD) | n |
|---|---|---|---|---|
| Water | 2019 | - 26.0 | 66.8 ± 2.2 | 4 |
| | | 3.1 | 59.1 ± 12 | 10 |
| | | 11.4 | 42.5 ± 5.3 | 10 |
| | | 20.5 | 37.2 ± 4.2 | 10 |
| | | 29.2 | 35.1 ± 4.9 | 10 |
| | | 37.5 | 36.2 ± 4.1 | 10 |
| | | 42.5 | 36.2 ± 5.4 | 6 |
| | 2020 | - 26.0 | 54.4 ± 2.1 | 3 |
| | | 3.1 | 63.3 ± 4.6 | 6 |
| | | 11.4 | 55.6 ± 3.4 | 6 |
| | | 20.5 | 53.6 ± 3.3 | 6 |
| | | 29.2 | 48.8 ± 2.3 | 5 |
| | | 37.5 | 50.7 ± 3.1 | 6 |
| | | 42.5 | 50.6 ± 2.4 | 3 |
| Woodchips | 2019 | 4.6 | 36.0 ± 5.1 | 9 |
| | | 19 | 27.4 ± 3.4 | 9 |
| | | 36.3 | 22.0 ± 1.0 | 9 |
| | 2020 | 4.6 | 49.5 ± 2.1 | 5 |
| | | 19 | 44.1 ± 2.2 | 6 |
| | | 36.3 | 34.0 ± 3.9 | 6 |
